# Accelerated Tempo of Cortical Neurogenesis in Down Syndrome

**DOI:** 10.1101/2025.09.29.679346

**Authors:** Jingwen W Ding, Chang N Kim, Marilyn R Steyert, Andrew T Yuan, David Shin, Dimitar Ivanov, Tomasz J Nowakowski, Alex A Pollen

**Affiliations:** The Eli and Edythe Broad Center of Regeneration Medicine and Stem Cell Research, University of California San Francisco, San Francisco, CA, USA; Department of Neurology, University of California San Francisco, San Francisco, CA, USA; Department of Neurological Surgery, University of California San Francisco, San Francisco, CA, USA; Department of Anatomy, University of California San Francisco, San Francisco, CA, USA; Department of Psychiatry and Behavioral Sciences, University of California San Francisco, San Francisco, CA, USA; Weill Institute for Neurosciences, University of California San Francisco, San Francisco, CA, USA

## Abstract

Down syndrome (DS), caused by trisomy 21 (TS21), is the most common genetic cause of intellectual disability^1,2^. The neurological impacts of DS first manifest during prenatal development through reduced radial glia (RG) neural stem cell proliferation, reduced cortical volume and imbalanced cortical cell types^3–6^. However, the developmental mechanisms underlying altered cortical neurogenesis in DS remain elusive. Here we show by high-throughput lineage tracing in organotypic culture that TS21 accelerates RG lineage progression, driving premature production of cortical inhibitory neurons (INs) and oligodendrocytes. Somatic lineage coupling connects dysregulated neurogenic tempo to altered cellular composition in the adult DS brain. Finally, lineage-resolved differential expression reveals elevated interferon responses specifically in RG biased to producing INs. Together, our findings link TS21 genomic abnormalities to candidate molecular pathways and developmental mechanisms altering the cellular landscape in DS with therapeutic relevance.

## Introduction

Individuals with Down syndrome (DS) exhibit disrupted neurodevelopment leading to lifelong learning, memory, and language impairment, as well as early-onset dementia^2,7–9^. Altered excitation/inhibition (E/I) ratio has been hypothesized as a candidate mechanism contributing to cognitive delay in individuals with DS in infancy and childhood^6,10,11^, but the underlying processes giving rise to this imbalance are currently unknown. One possible mechanism involves increased abundance of inhibitory neurons (INs) generated during development^12–14^. In mice, most cortical INs arise from progenitors in the ganglionic eminences (GE)^15^, but human cortical radial glia (RG) can also produce cortical-like INs at late stages of neurogenesis^16,17^. While recent studies have begun to characterize the molecular and cellular consequences of TS21 during cortical development^18–20^, the dynamic processes of RG differentiation, maturation, and lineage progression have been difficult to assess in a human context. By performing high-throughput lineage tracing in human primary cortical organotypic culture, we identify a dysregulated tempo of cortical neurogenesis in TS21, characterized by precocious production of cortical INs and linked to elevated interferon (IFN) signaling.

## Results

### Single cell lineage tracing in the TS21 developing cortex

To systematically assess progenitor cell phenotypes in the TS21 cortex, including proliferative capacity and cell fate biases, we generated a lineage-resolved single cell RNA sequencing (scRNA-seq) dataset from a cohort of 14 individuals (CTRL, n=7; TS21, n=7) spanning middle and late stages of cortical neurogenesis and early gliogenesis (Gestational week (GW) 14-23.5) (Table S1). We applied STICR, a recently established tool for systematic clonal cell lineage tracing through lentivirally delivered static barcodes^16^ (Figure 1A). To enrich for neural progenitors, we locally transduced germinal zones (GZ) on *ex vivo* organotypic slice culture and isolated green fluorescent protein (GFP) positive cells after 13-14 days for scRNA-seq to recover transcriptomic cell identities and lineage barcodes (Figure 1A). This yielded 147,861 single cells passing quality control criteria across all samples, including 64,702 cells from CTRL and 83,159 from TS21 samples (Figures 1B and S1A-C). We further performed clone assignment based on lineage barcodes and considered multicellular clones containing at least 3 cells, recovering 6,100 clones (CTRL, 2,453; TS21, 3,647) representing newborn populations following STICR labeling that reflect differentiation dynamics of cortical RG (Figures 1C-E and S1D-G).

**Figure 1:**
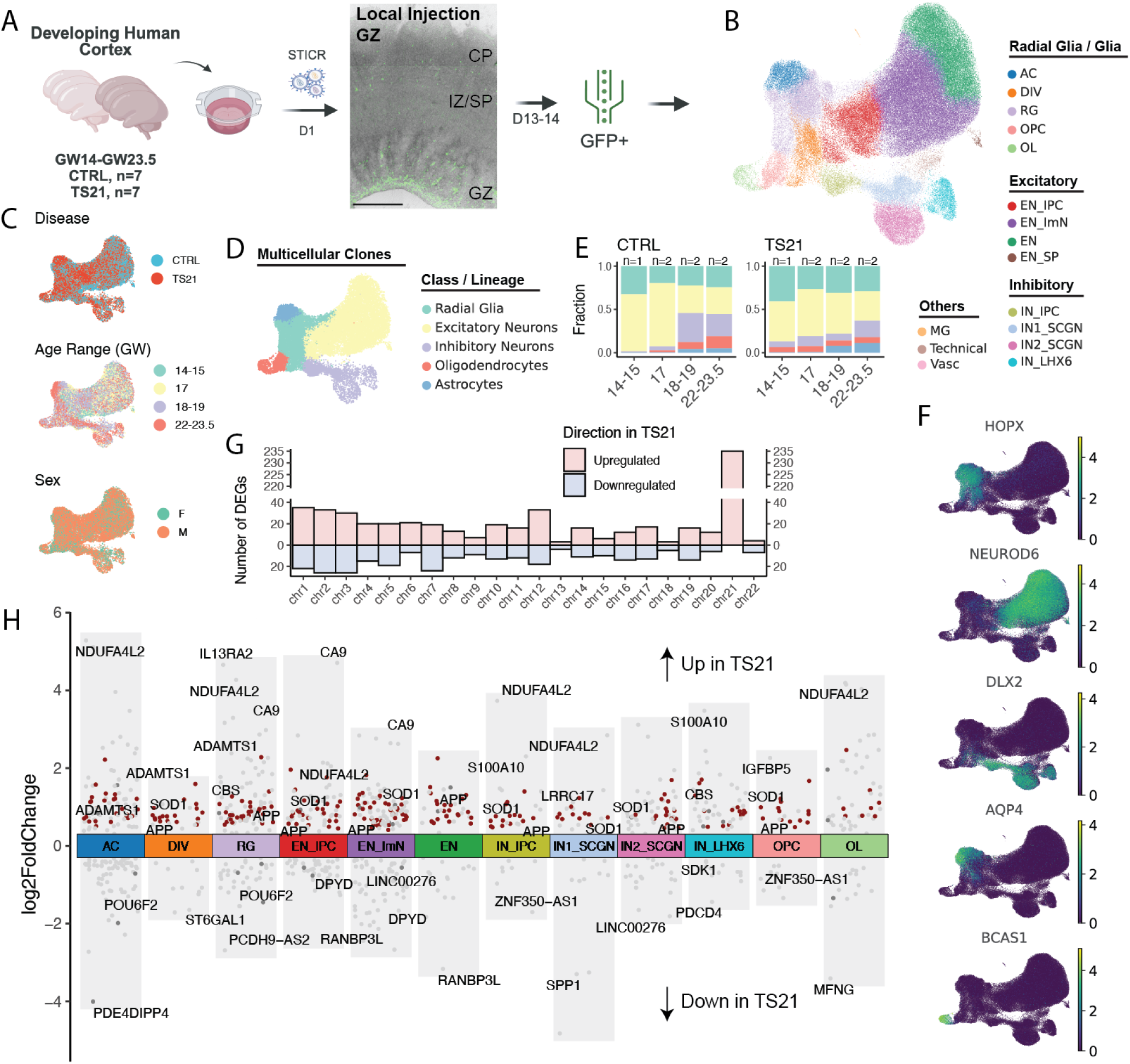
Single cell lineage tracing in the developing cortex in TS21. A. Experimental design for high-throughput single cell lineage tracing on organotypic slice culture of the developing cortex in TS21, with an example image to show local GFP labeling of neural progenitor cells at GZ on infection day 5. B. UMAPs of cells collected on day 13-14 post local transduction, colored by supervised cell types. C. UMAPs of cells in multicellular clones (more than 2 cells per clone), colored by disease status, age range and sex (Table S1). D. UMAP showing cells in multicellular clones, colored by cell class. E. Stacked barplots showing distributions of cell lineage at each age range in multicellular clones. F. UMAPs colored by the expression of *HOPX, NEUROD6, DLX2, AQP4* and *BCAS1*. G. Barplot showing numbers of up- and down-regulated DEGs in TS21, grouped by chromosome. H. Volcano plot for DEGs in the cell type level between CTRL and TS21. DEGs with an absolute log_2_FC change more than 0.2 were shown. DEGs on HSA21 were highlighted in red. Scale bar: 500 μm. GW, gestational week; CTRL, control; TS21, trisomy 21; GZ, germinal zone; IZ, Intermediate zone; SP, subplate; CP, cortical plate; GFP, green fluorescent protein; RG, radial glia; AC, astrocytes; EN, excitatory neuron; IN, inhibitory neuron; IPC, intermediate progenitor cell; ImN, immature neuron; DIV, dividing; OPC, Oligodendrocyte progenitor cell; OL, oligodendrocyte; MG, microglia; Vasc, vascular; DEGs, differentially expressed genes; log_2_FC, log2 fold change; HSA21, human chromosome 21

scRNA-seq recovered four principal cortical cell lineage trajectories - excitatory (ENs) and inhibitory neurogenesis (INs) and gliogenic trajectories including astrogenesis (AC) and oligogenesis (OPC and OL), marked by *NEUROD6*, *DLX2, AQP4 and BCAS1*, respectively (Figures 1F and S1A-B). Integrating all samples from both conditions revealed consistent representation of cells derived from independent biological replicates across cell types with limited batch effects (Figures 1C and S1C). Reference mapping to a developmental cortical cell atlas^17^ revealed representation of *in vivo* cell types with strong transcriptional correlations (Figures S1H-I), confirming that the *ex vivo* slice culture system recapitulated *in vivo*-like gene expression, cell types, states, and differentiation dynamics. Most neurons displayed immature profiles, consistent with their recent derivation from RG. As previously described^16,17^, we observed sequential stage-dependent changes in cell type composition along midgestation, with ENs replaced by INs, AC and OPC/OL, reflecting a cell fate switch of RG from producing EN to IN and glia cell types (Figure 1E). In TS21, we observed an increased number of IN progenitors (IN_IPCs), consistent with a recent study of post-mortem tissue^19^. In addition, we detected cortical IN and OPC/OL at earlier stages in TS21, indicating potential dysregulation of RG differentiation and fate commitment (Figure 1E).

Differential gene expression analysis revealed global upregulation of genes on human chromosome 21 (HSA21) (Figures 1G-H and S2A; Table S2). These include *APP* and *SOD1*, genes that have been implicated in neurodegenerative disorders including Alzheimer’s disease^3^ and amyotrophic lateral sclerosis (ALS)^21^; *NDUFA4L2* and *CBS*^22^, genes essential to mitochondria function (Figure 1H). Changes in gene expression on other chromosomes were also observed, indicating transcriptome-wide gene dysregulation in TS21. Pathway analysis highlighted impaired protein translation and processing, inflammatory response, glycolysis, and oxidative stress in TS21 (Figures S2B and S2C), consistent with previous findings^23–26^.

### Dysregulated neurogenic tempo in TS21

The changes in relative abundance of ENs, INs and glia raised the possibility that TS21 alters RG cell fate choices and developmental tempo. Lineage tracing revealed that the most abundant multiclass clones contained RG and ENs in both conditions, but the overall proportion of RG and EN-containing clones decreased in TS21 (CTRL, ∼40%; TS21, ∼25%), replaced by clones containing glia (Figures 2A and S3A). This observation was further supported by lineage coupling scores quantified as the normalized barcode covariance between cell types^27^ (Figure S3B), suggesting increased gliogenesis in TS21 and consistent with previous reports^4,20,28^.

**Figure 2:**
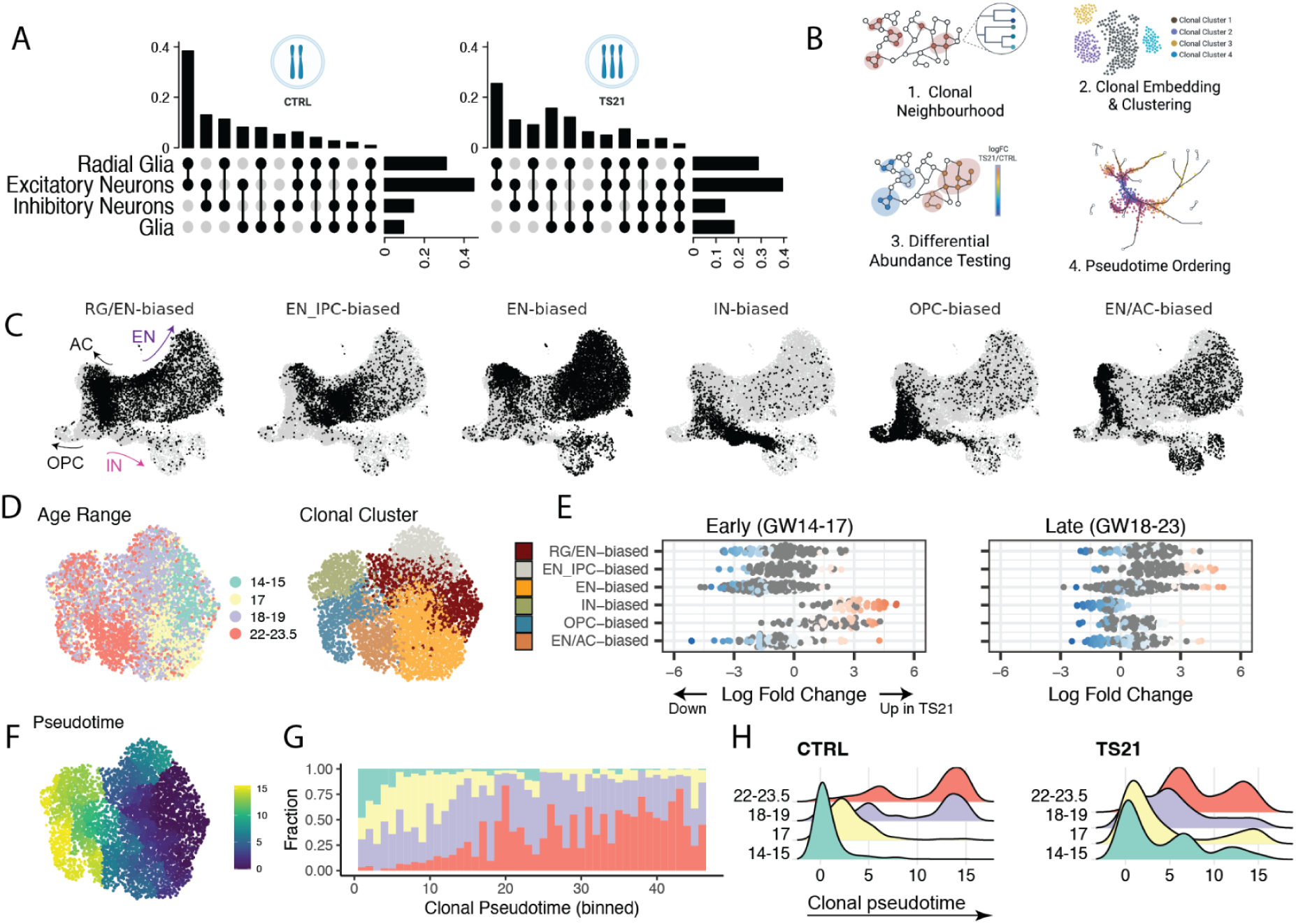
Dysregulated neurogenic tempo in TS21. A. Upset plots of cell class compositions in multicellular clones that contain cells from more than one class (multi-class clones) in CTRL (left) and TS21 (right). B. Schematics for clonal analysis pipeline. C. UMAPs of cells for cell type distributions in clonal clusters identified using scLiTr^52^: RG/EN-, EN_IPC-, EN-, IN-, OPC- and EN/AC-biased clusters. D. UMAPs of clones, colored by age range and clonal clusters. E. Beeswarm plots showing differential abundance between CTRL and TS21 in neighborhoods of clones in each clonal cluster. Samples were divided into two age groups that correspond to before and after the beginning of IN production in CTRL: early, GW14-17 (left) and late, GW18-23 (right). Neighborhoods that had significant changes (FDR=0.05) between disease and age conditions were colored by their log_2_FC. F. UMAPs of clones colored by pseudotime, inferred using Monocle3^53^ based on composition proximity of clones. G. Stacked barplots showing fractions of clones in each age range along binned clonal pseudotime. The x-axis represents bin numbers along pseudotime. Color denotes age range defined in panel D. H. Ridge plots of pseudotime distribution in clones grouped by age in CTRL (left) and TS21 (right).

To further investigate developmental mechanisms underlying composition changes, we grouped 6,100 clones into 6 clusters based on their fate outcomes using scLiTr^29^, a tool that embeds clones into a low-dimensional space and identifies major groups and trajectories. We identified RG/EN-, EN_IPC-, EN-, IN-, OPC- and EN/AC-biased clones and compared differential abundance of clonal clusters between stages and disease conditions (Figures 2C-D and S3C-D). Across developmental stages, we observed sequential emergence of IN-, OPC- and AC-biased clones in CTRL, as expected^17,30^ (Figure S3E). However in TS21 samples, we observed premature emergence of IN- and OPC-biased clones at early stages (GW14-17) at the expense of clones favoring the excitatory lineage (Figure S3E), consistent with observations at the composition level.

As cortical inhibitory neurogenesis is normally restricted to late midgestation^16,17^, we divided samples into two age groups: early (GW14-17, CTRL: n=3; TS21: n=3) and late (GW18-23, CTRL: n=4; TS21: n=4) midgestation to examine stage-dependent alterations in RG lineage in TS21. To robustly identify differential abundance at the clone level, we further applied a cluster-free approach to sensitively detect differential abundance independent of discrete cluster annotation by grouping clones with similar fate outcomes into overlapping neighborhoods^31^ (Figure S3F). Contrasting between disease and age groups again revealed significant enrichment of IN- and OPC-biased clones specifically at early ages of TS21 at the expense of other classes of EN-biased clones (Figure 2E). Interestingly, opposing phenotypes were observed at later stages, where depletion of IN- and OPC-biased clones in TS21 was observed with an overall preference to classes of EN-biased clones in comparison to CTRL (Figures 2E and S3F). Similarly, composition analysis in cells from multicellular clones showed consistent results of enrichment in *SCGN* expressing INs in early TS21 samples, reinforcing early acceleration of RG fate commitment in TS21 (Figures S3G-S3I).

To further characterize this change in RG lineage progression, we constructed a pseudotime trajectory at the clone level (Figure 2F). The inferred clonal pseudotime reproduced the progression in cell fate choices across stages (Figure 2G). Comparing the pseudotime distribution between disease conditions and age groups revealed acceleration at GW14-17 but delay at GW18-23 in RG lineage progression in TS21, indicating dysregulated neurogenic tempo in TS21 (Figure 2H).

### Early emergence of cortical INs in the DS brain

To investigate the physiological relevance of accelerated IN production observed by *ex vivo* lineage tracing, we quantified EN and IN populations using immunohistochemistry (IHC) from fixed tissue (Figures 3A and 3B). To minimize technical biases, we applied an automated scalable pipeline using Cellpose^32^ for signal segmentation and quantification (Methods). Across a cohort of 21 individuals (Table S3), we observed a significant increase of SCGN expressing IN populations in early TS21 samples (GW14-18, CTRL, n=5; TS21, n=5) in both GZ (Two-sided Wilcoxon test, p=0.0079), where newborn populations reside, and intermediate zone (IZ) and subplate (SP) (p=0.032), where immature populations migrate through before they reach cortical plate (CP) (Figures 3C and S4). While we did not observe significant changes in the NEUROD2 positive EN populations, we noted an increase of the overall SCGN to NEUROD2 ratio at early stages (Figure 3C). These results further indicate a transient increase in inhibitory neurogenesis at early midgestation in TS21 *in vivo*.

**Figure 3:**
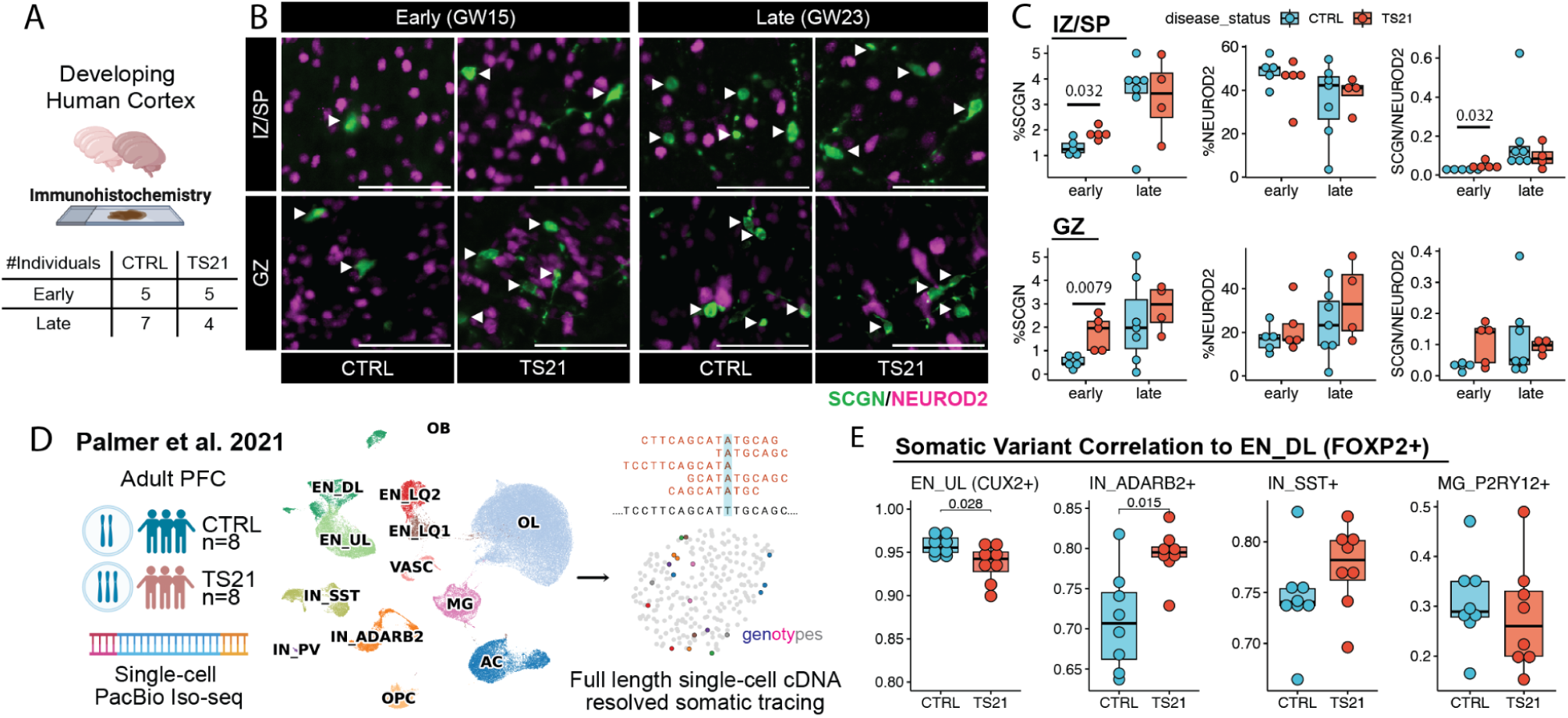
Early emergence of cortical INs in the TS21 brain *in vivo*. A. Experimental design of *in vivo* quantification of cell type markers through IHC in CTRL and TS21. A cohort of 21 individuals were included for quantification and divided into two age groups: early, GW14-18 and late, GW19-23.5. B. Representative images of immunohistochemical labeling of SCGN and NEUROD2 at GZ and IZ/SP in samples aged GW15 and GW23, representative of early and late age groups. C. Barplot showing fractions of SCGN and NEUROD2 expressing populations in GZ (top) and IZ/SP (bottom), grouped by disease status and age groups. Each dot represents an independent individual. D. Schematics for cell-type-resolved lineage coupling analysis based on somatic mosaicism in the adult DS PFC using single cell Iso-Seq libraries described in Palmer et al., 2021^12^. A cohort of 16 individuals were included for this analysis. E. Barplots showing Pearson correlation coefficients of somatic variants between *FOXP2*+ DL ENs and other cell types: *CUX2*+ UL ENs, *ADARB2*+ and *LHX6*+ INs and *P2RY12*+ MG. Scale bar: 50 μm. IHC, immunohistochemistry; PFC, prefrontal cortex

We next asked if the effects of altered neurogenic tempo persist into the adult DS brain. We reasoned that precocious production of cortical INs could be reflected by altered lineage coupling in the adult brain. The shift to cortical IN production normally occurs late, following upper layer (UL) EN production^16,17^. However, we hypothesized that earlier IN production in DS could enhance IN lineage coupling to earlier born deep layer (DL) ENs. To test this prediction, we analyzed single cell long read Iso-seq libraries generated in the adult DS prefrontal cortex (PFC)^12^ and performed somatic mosaicism informed lineage tracing with cell type resolution (Figure 3D, Methods). We re-processed the data and batch-corrected representation of 16 individuals across all cell types (Figures S5A and S5B). We focused on major cell types that are well represented in the PFC (Figure S5C), including ENs (*FOXP2*+ DL and *CUX2*+ UL) that are generated sequentially from cortex and INs (*ADARB2*+ and *SST*+) that have dual origins from both cortex and caudal and medial GE (CGE and MGE)^33,34^, respectively. *ADARB2*+ INs transcriptionally correspond to *SCGN*+ INs in the developing cortex while *SST*+ INs represent *LHX6*+ INs (Figures S5C and S5D). We also included *P2RY12*+ microglia as a control cell type that diverge early from other neuronal populations. Utilizing the increased genome coverage of full-length libraries we were able to generate genotypes with cell type resolution. Comparison of somatic variant profiles revealed a significant increase in Pearson correlation between DL ENs and *ADARB2*+ INs in DS, suggesting a closer clonal linkage between the two populations, consistent with early emergence of these IN populations (CTRL, n=8; TS21, n=8, Two-sided Wilcoxon test, p=0.015) (Figure 3E). While it is difficult to estimate the contributions of CGE or locally derived populations of *ADARB2+* INs due to their mixed origins, this change in clonal linkages further supports precocious cortical IN production in DS. Together, these results suggest an early acceleration of cortical RG lineage resulting in premature emergence of cortical-like INs prenatally that can persist into adulthood.

### Lineage-specific upregulation of IFN response

We were motivated to further investigate the molecular mechanisms underlying precocious cortical IN production in TS21 and used DEseq2^35^ to identify differentially expressed genes in the radial glia class (RG, DIV) of each clonal cluster between disease conditions. This lineage-resolved analysis revealed previously characterized genes that have been implicated in RG fate decisions (Table S4). For example, upregulation of IFN stimulated genes was observed in AC-biased clones, consistent with the known role of its downstream pathway JAK-STAT in promoting neurogenic to gliogenic switch in RG^3,36^ (Figures 4A and S6A-B). Focusing on the IN and EN lineage, pathway analysis revealed opposing changes in JAK-STAT signaling pathway, with significant upregulation in IN-biased RG but downregulation in EN-biased RG in TS21 (Figures 4B and 4C), suggesting a potential role of IFN in excitatory to inhibitory neurogenic switch.

**Figure 4:**
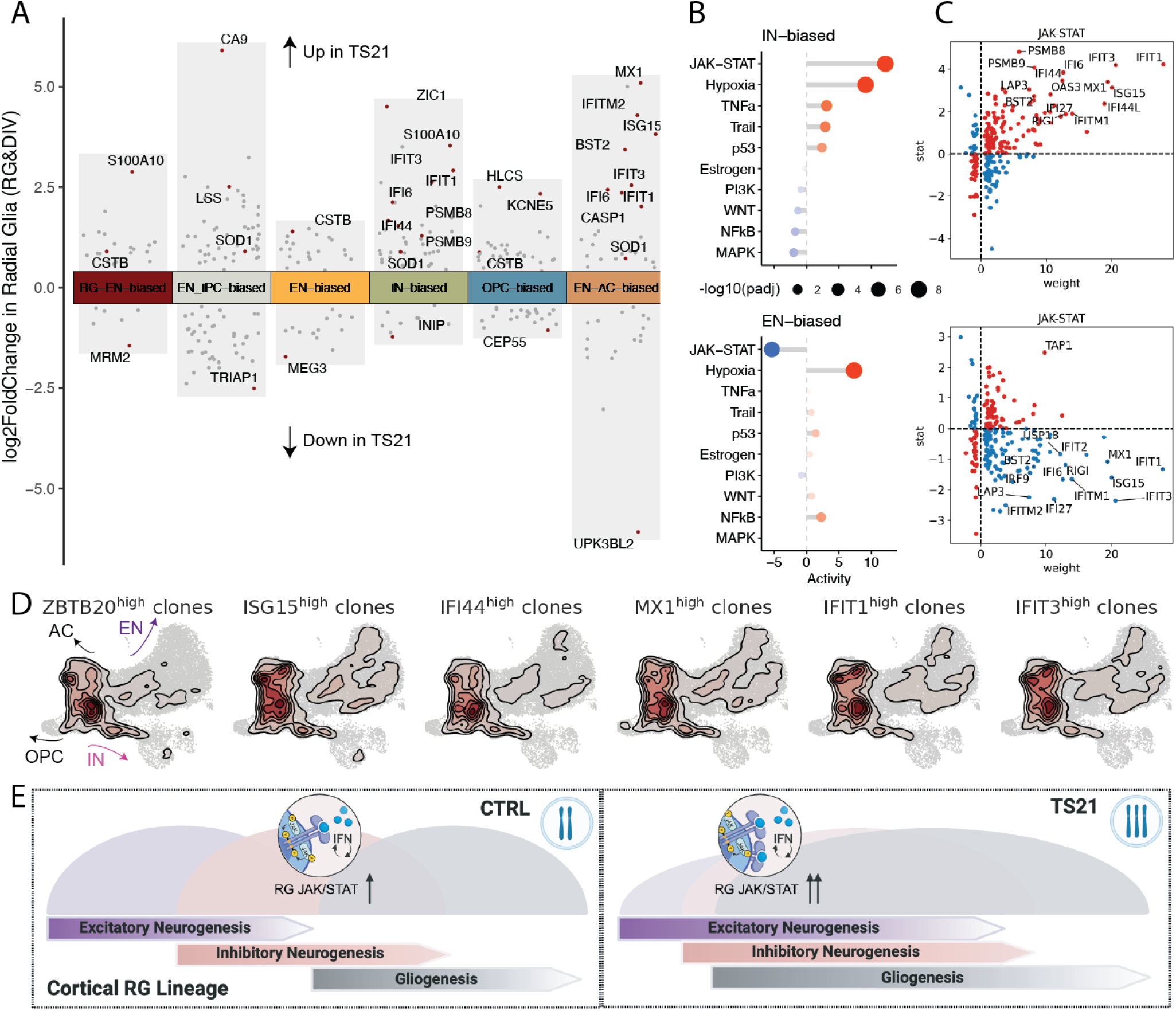
Elevated IFN response in TS21 in cortical inhibitory neurogenesis. A. Volcano plot for DEGs between CTRL and TS21 in the radial glia class (RG, DIV) grouped by clonal clusters. DEGs with an absolute log_2_FC change more than 0.2 were shown. Top DEGs in each clonal cluster and IFN stimulated genes were highlighted and labeled. B. Cleveland’s dot plots showing pathway activity in DEGs in IN- (top) and EN- (bottom) biased RG. Size denotes -log_2_(adjusted p value) and color denotes pathway activity inferred using a univariate linear model through PROGENy^54^. C. Scatterplots of genes associated with the JAK-STAT pathway in IN- (top) and EN- (bottom) biased RG. The x-axis denotes the weight of the target gene in the pathway obtained from PROGENy^54^. The y-axis represents the Wald test statistic from DESeq2^35^, defined as the estimated log_2_FC divided by its standard error. D. Kernel density estimate (KDE) plots of cells from top 15% clones where the radial glia class highly expresses IFN responsive genes: *ZBTB20*, *ISG15*, *IFI44*, *MX1*, *IFIT1* and *IFIT3*. E. Graphic summary of findings: cortical RG in TS21 display premature inhibitory neurogenesis at the expense of ENs and dysregulated developmental tempo, associated with elevated IFN response and activation of JAK-STAT signaling pathway. IFN, interferon

Four genes encoding receptors for IFNs (*IFNAR1*, *IFNAR2*, *IFNGR2* and *IL10RB*) are located on HSA21 and are triplicated in DS, suggesting potential dysregulation of IFN response in the DS brain as observed in other tissues^37–41^. Indeed, strong upregulation of these four IFN receptors, as well as IFN stimulated genes, was observed in dividing progenitors in the developing TS21 cortex (Figures S6C and S6D). Individuals with DS regression disorder (DSRD), a condition that occurs in a subset of DS patients characterized by sudden loss of neurological function, exhibit radiographic features associated with increased IFN activation in the bilateral basal ganglia^42^. Normalizing the copy number of the IFN receptor locus attenuates the developmental delay and cognitive deficits in a DS mouse model^43^, further nominating elevated IFN response and downstream JAK-STAT activation as candidate mechanisms underlying neurodevelopmental phenotypes in DS. Consistent with results from differential gene expression analysis, the expression of IFN stimulated genes in RG increases along clonal pseudotime, which coincides with inhibitory neurogenesis (Figure S6E). High expression of IFN stimulated genes individually and high activity of IFN and JAK-STAT pathway in RG marks clones that are biased towards IN and AC fates (Figures 4D and S6F), linking IFN activation to RG fate consequence. Collectively, our results suggest a role of elevated IFN response in regulating EN/IN production through altering RG lineage progression (Figure 4E).

## Discussion

E/I imbalance has emerged as a prevailing candidate mechanism underlying cognitive deficits in DS^6,10^, supported by overrepresentation of a subset of cortical INs in the adult DS brain^12^. Our lineage-resolved analysis of RG cell fate in human organotypic slice culture suggests accelerated maturation of RG in TS21, including precocious inhibitory neurogenesis. The model of accelerated lineage progression of RG is reinforced by somatic lineage tracing in the adult DS brain, where a significant increase of somatic variant correlation of DL ENs with *ADARB2*+ INs indicates the early cortical origin of these INs, linking the dysregulated neurogenic tempo to altered cellular compositions in the adult brain. The transient increase of IN-biased clones is partially offset through enhanced EN production at late midgestation, but the accelerated timing of IN production also impacts the abundance of other RG progenies. A postnatal atlas of the DS prefrontal cortex identified increased UL ENs at the expense of DL ENs^20^, a phenotype that could arise, in part, from precocious local IN generation skewing EN subtype balance.

Multiple HSA21-encoded genes, including *DYRK1A*^3,44^, *APP*^45^, *OLIG1*^13^, and *OLIG2*^46^, have been implicated in RG proliferation, differentiation and fate commitment. Four IFN receptors are encoded by HSA21 and overexpressed in DS^37^. Three types of IFNs are involved as ligands for these receptors, but they convergently activate the downstream JAK-STAT pathway, which has been hypothesized to promote RG fate switch from neurogenesis to gliogenesis in TS21^4,47^. Our finding of lineage-specific upregulation of IFN responses in IN-biased RG suggests a role of JAK-STAT signaling in TS21 in promoting RG lineage transition from EN to IN production. This model is consistent with the role of LIF signaling, upstream of JAK-STAT, in promoting IN production in forebrain organoids^48^. JAK inhibition (JAKi) has shown efficacy in improving diverse immune skin pathologies in DS^38,49,50^. Interestingly, cooccurance of neurological conditions-including seizure and movement disorders and structural brain abnormalities-along with autoantibody positivity in DS patients^49^ has been noted, supporting dual roles of elevated IFN response in disrupting neural development and increasing autoimmune burden^51^. Together, our study reveals developmental mechanisms contributing to altered cellular landscape in DS and provides candidate molecular pathways with therapeutic relevance.

## Resources

### Lead contact

Further information and requests for resources and reagents should be directed to and will be fulfilled by the lead contact, Alex A. Pollen.

### Materials availability

This study did not generate new unique reagents.

### Data and code availability

- Raw sequencing data will be made available through dbGaP upon publication.
- Any additional information required to reanalyze the data reported in this paper is available from the lead contact upon request.

## Acknowledgments

The authors thank A. Molofsky, S. Anderson, Pollen and Nowakowski lab members for valuable comments, J. Srivastava for assistance performing cell sorting at the Gladstone Flow Cytometry Core, supported by NIH S10 RR028962 and S. Wang for transferring samples. Sequencing was performed at the UCSF CAT, supported by UCSF PBBR, RRP IMIA, and NIH 1S10OD028511-01 grants. Figures were made, in part, with BioRender. This work was supported by the following funding sources: CIRM Fellowship (J.W.D.), Discovery Fellowship (C.N.K.), National Institutes of Health R01MH134981-01, DP2MH122400-01 (A.A.P.), UM1MH130991 (A.A.P., T.J.N.), R01NS123263 (T.J.N.), Schmidt Futures Foundation (A.A.P., T.J.N.), William K. Bowes Jr. Foundation (A.A.P. and T.J.N.). Shurl and Kay Curci Foundation (A.A.P., T.J.N.), the California Institute for Regenerative Medicine (CIRM) DISC0-14429 (T.J.N.), DISC4-16285 (A.A.P), as well as gifts from Esther A. & Joseph Klingenstein Fund (T.J.N.) and the Sontag Foundation Distinguished Scientist Award (T.J.N.). T.J.N. is a New York Stem Cell Foundation Robertson Neuroscience Investigator. A.A.P. is a New York Stem Cell Foundation Robertson Investigator.

## Author Contributions

Conceptualization: J.W.D., C.N.K., T.J.N., A.A.P.

Methodology:J.W.D., C.N.K., M.R.S., A.T.Y., D.S., D.I.

Investigation: J.W.D., C.N.K.

Visualization: J.W.D., C.N.K.

Funding acquisition: T.J.N., A.A.P.

Project administration: T.J.N., A.A.P.

Supervision: T.J.N., A.A.P.

Writing – original draft: J.W.D., C.N.K., T.J.N., A.A.P.

Writing – review & editing: J.W.D., C.N.K., T.J.N., A.A.P.

## Declaration of interests

The authors declare no competing interests.

## Supplemental Figures

**Figure S1:**
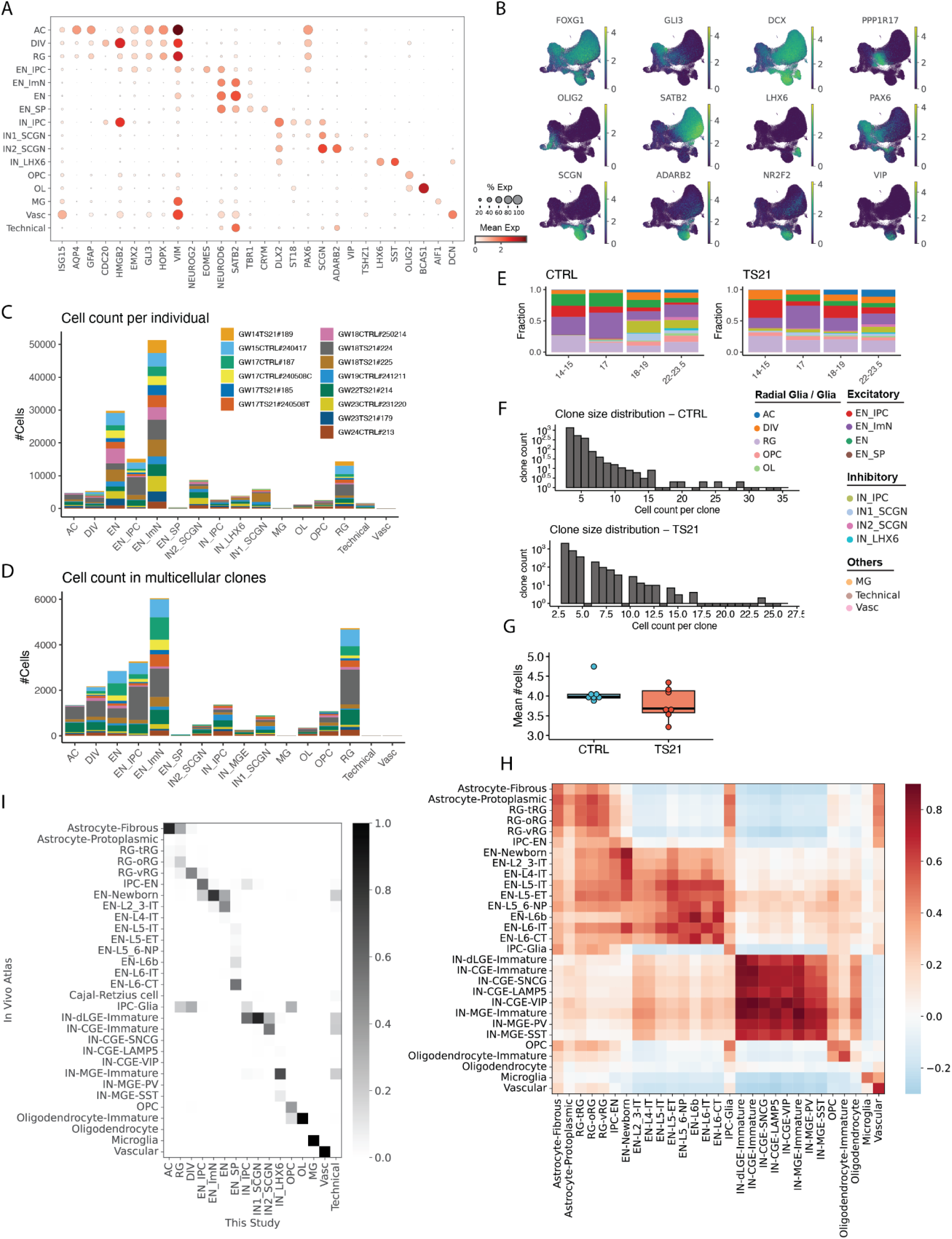
Single cell lineage tracing and comparison to *in vivo* atlas, related to Figure 1. A. Dotplot of the expression of cell type markers for the supervised cell types. The size of each dot denotes the fractions of cells in the group where the gene is expressed and the color denotes mean gene expression in the group. B. UMAP of cells colored by the expression of cell type markers. C. Stacked barplots showing cell counts of individuals in each cell type. D. Stacked barplots showing cell counts of individuals in each cell type in multicellular clones. E. Stacked barplots showing fractions of cell types between ages in multicellular clones. F. Histograms for distributions of clone size in CTRL (top) and TS21 (bottom). Clones with fewer than three cells were removed from downstream analyses. G. Boxplot for mean clone size in each individual in CTRL and TS21, with each dot shows value from one independent individual. H. Pearson correlation coefficients of top 25 cell marker gene expression defined in Wang et al^17^ *in vivo* atlas to data obtained in this study. Cell type annotation was done using reference mapping to the *in vivo* atlas. I. Heatmap for fraction overlap of labels transferred from reference cell types^17^ to the cell types defined in this study.

**Figure S2:**
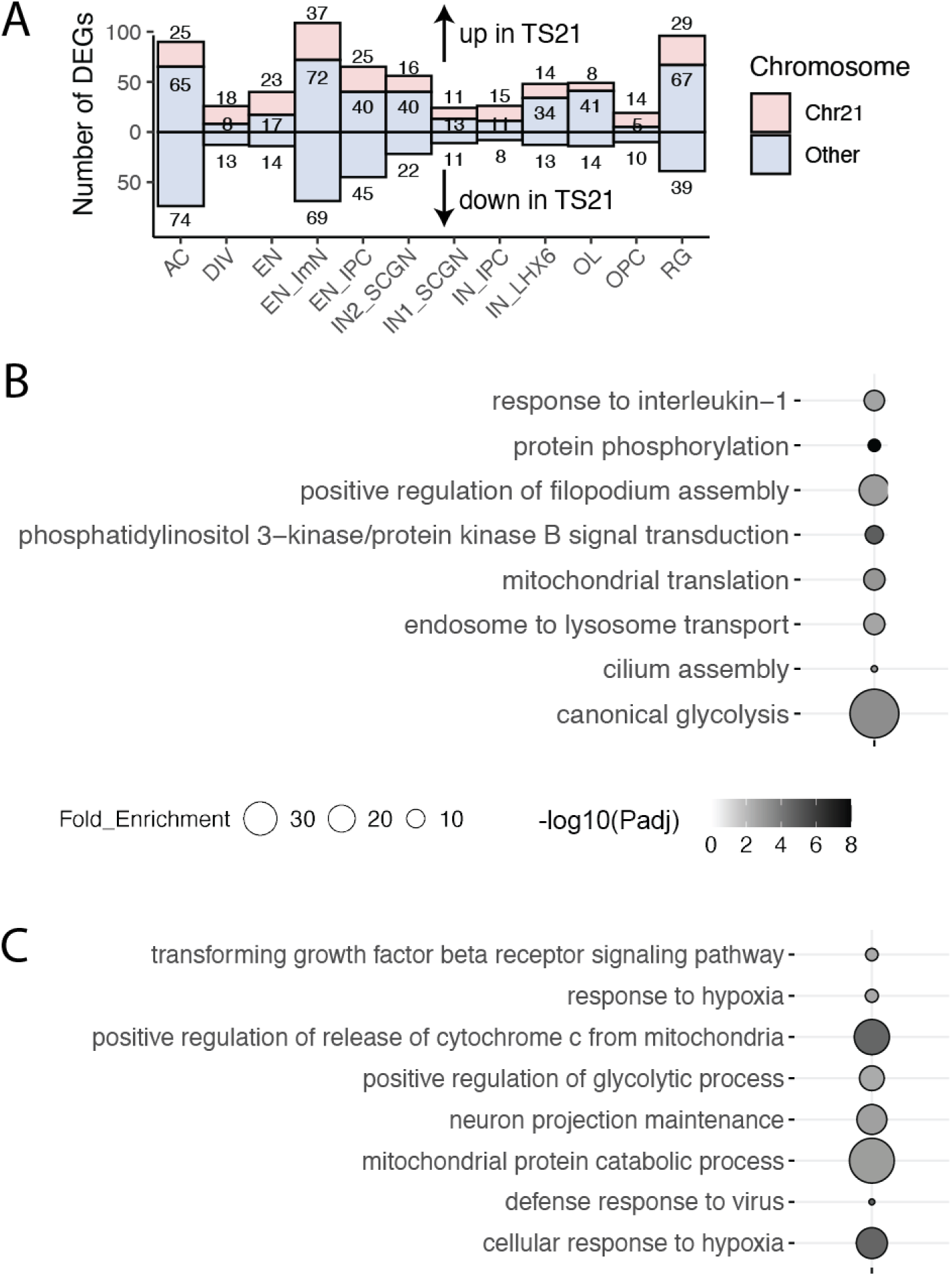
Differential gene expression analysis between CTRL and TS21, related to Figure 1. A. Barplot showing numbers of up- and down-regulated DEGs in TS21, grouped by cell types. B. Enrichment plot for GO terms in biological processes enriched in DEGs identified in all cell types in TS21 using pathfindR^55^. Color denotes -log_10_(adjusted p value) and dot size denotes fold enrichment in each category. C. Enrichment plot for GO terms in biological processes enriched in DEGs identified in the radial glia class (RG, DIV) in TS21. GO, gene ontology

**Figure S3:**
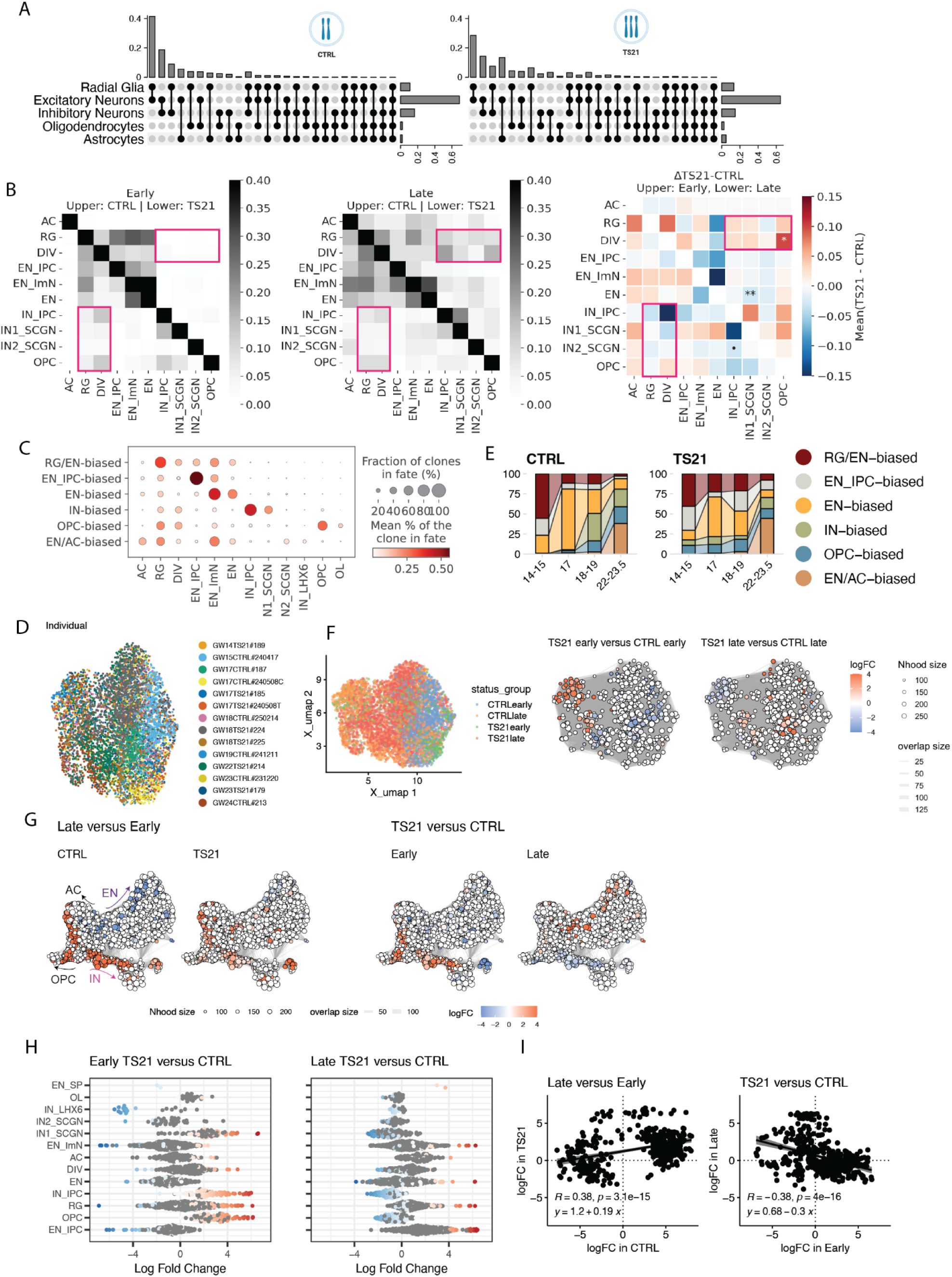
Clonal clustering and differential abundance testing, related to Figure 2. A. Upset plots of cell class compositions in multi-class clones in CTRL (left) and TS21 (right), with separate rows for astrocyte and oligodendrocyte lineage. B. Heatmaps for lineage coupling score matrix in major cortical derived cell types in early (left) and late (middle) midgestation, with upper right showing TS21 and lower left showing CTRL. (right) Heatmap for changes in lineage coupling score matrix in TS21 compared to CTRL, with upper right showing early and lower left showing late midgestation. Only major cell types that were robustly detected in all samples were included. Red boxes highlight changes in lineage coupling between RG progenitors and INs and OPCs in early TS21 samples. C. Dotplot for distributions of supervised cell types in each clonal cluster identified using scLiTr^52^: RG/EN-, EN_IPC-, EN-, IN-, OPC- and EN/AC-biased clusters. The size of each dot denotes the fractions of clones in each clonal cluster where the cell type is detected and the color denotes the mean percentage of each cell type in clones. D. UMAPs of clones colored by individuals. E. Stacked barplot of distributions of clonal cluster fractions along ages in CTRL and TS21, colored by clonal clusters. F. UMAPs for differential abundance testing in clones between disease conditions and age groups using Milo. (left) UMAPs of clones colored by conditions tested; (middle-right) Neighborhood graphs of differential abundance testing colored by log_2_FC (FDR=0.05) comparing CTRL and TS21 in early (middle) and late (right) age groups. Red represents enrichment in TS21. Opposite composition changes at early and late stages were noted. G. UMAPs for differential abundance testing in cells contrasting between age groups (left) and disease status (right). Neighborhood graphs of differential abundance testing colored by log_2_FC (FDR=0.05). Red represents enrichment in late (left) and in TS21 (right), respectively. H. Beeswarm plots for differential abundance results shown in the right panel in Figure S3G grouped by cell type in early (left) and late (right) age groups. Neighborhoods that had significant changes between disease and age conditions were colored by their log_2_FC. I. Scatterplots showing log_2_FC in each neighbourhood (left) contrasting early and late groups in CTRL (x-axis) and TS21 (y-axis) and (right) contrasting CTRL and TS21 in early (x-axis) and late (y-axis) groups. Pearson correlation coefficients, p values and with linear regression lines and equations were labeled.

**Figure S4:**
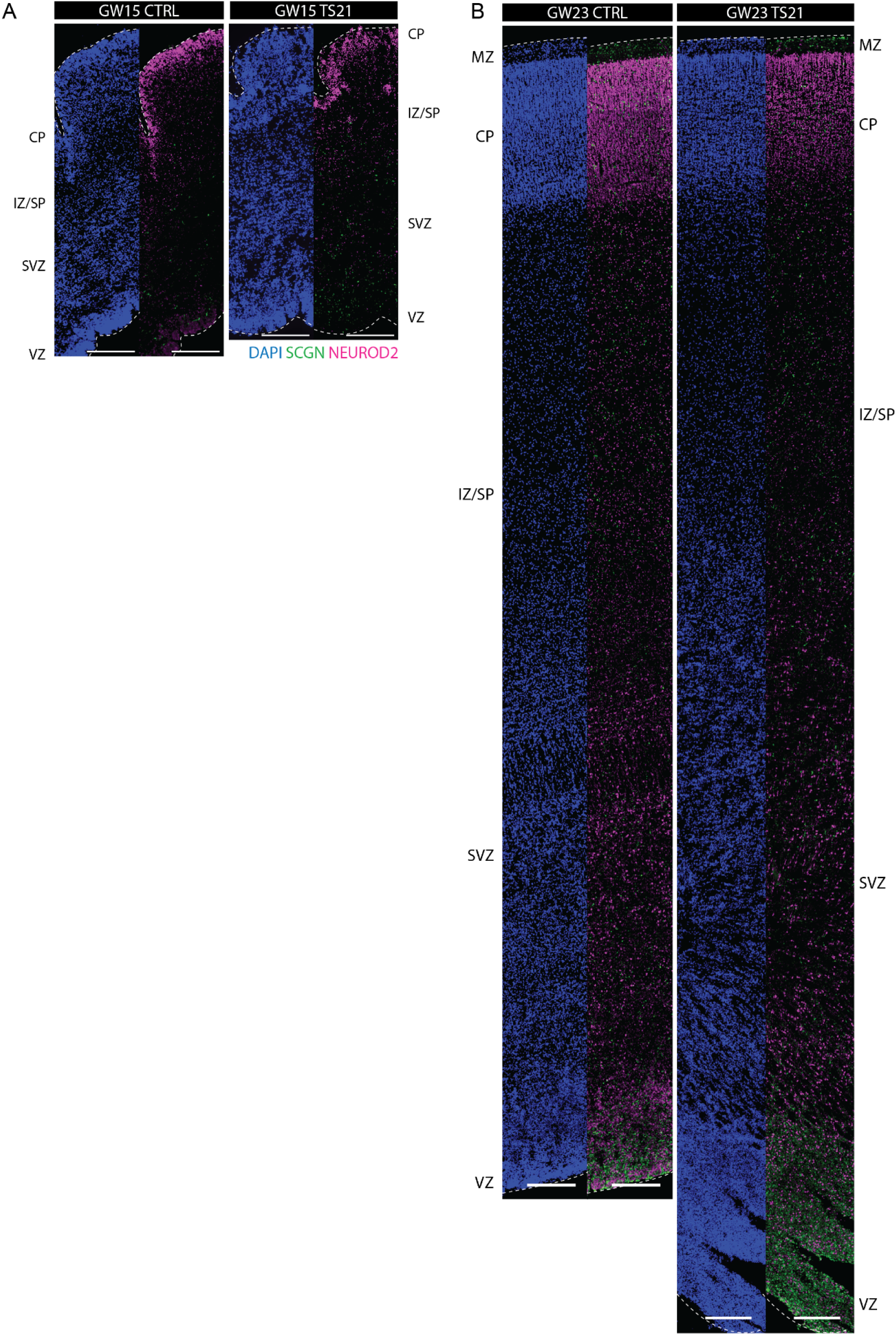
*In vivo* IHC quantification of cortical INs and ENs, related to Figure 3. A. Tile scans of immunohistochemical labeling of SCGN and NEUROD2 in GW15 CTRL and TS21 samples. B. Tile scans of immunohistochemical labeling in GW23 samples. Scale bar: 250 μm. VZ, ventricular zone; SVZ, subventricular zone; MZ, marginal zone

**Figure S5:**
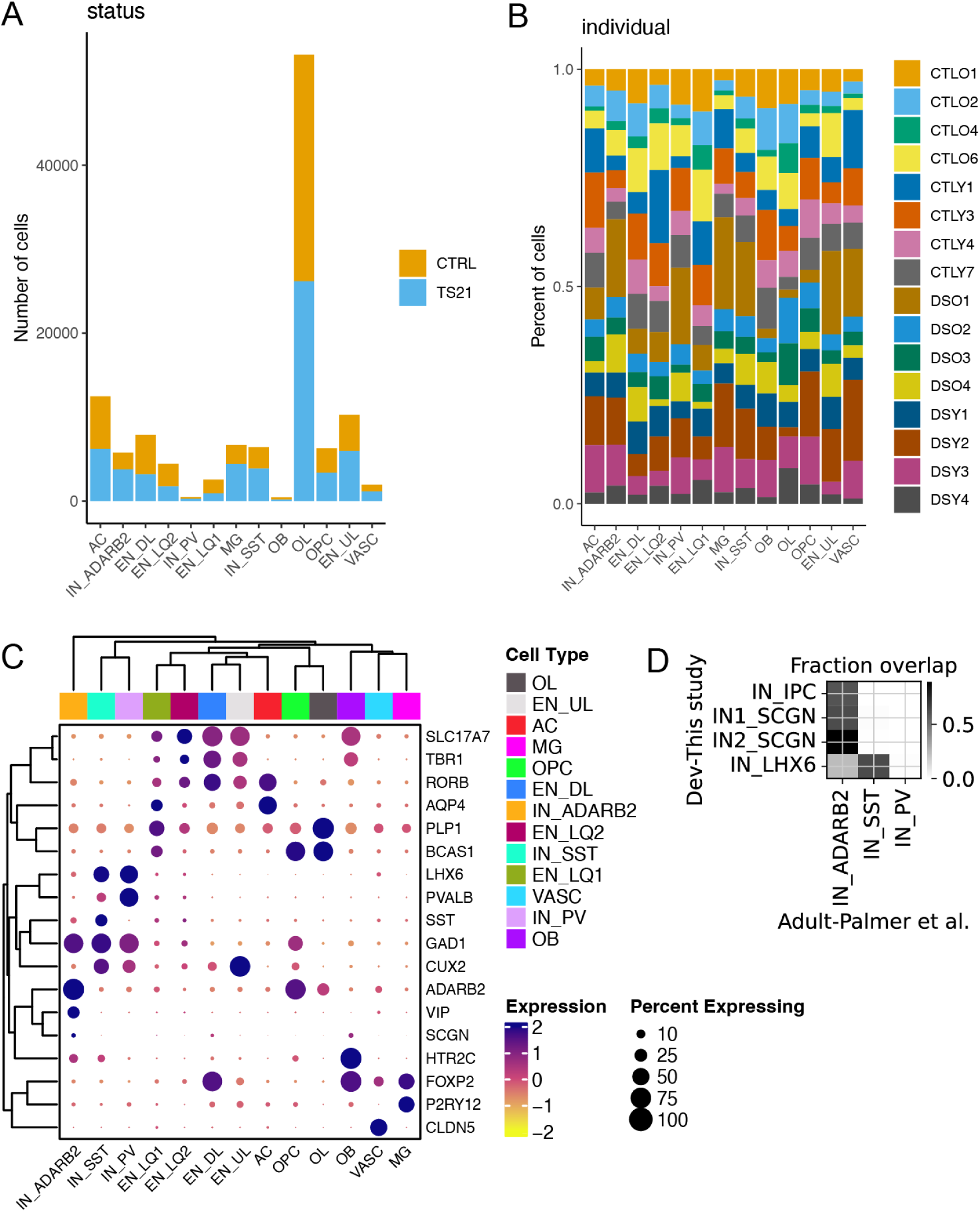
Analysis of scRNA-seq in the postmortem adult PFC samples, related to Figure 3. A. Stacked barplot showing cell counts in CTRL and TS21 across cell types. Data obtained from Palmer et al., 2021^12^. B. Stacked barplot showing cell fractions of individuals across cell types. C. Dotplot of marker gene expression across cell types. Note that *ADARB2*+ INs also co-express *SCGN* and *VIP*. D. Heatmap for fraction overlap of IN labels transferred from the Palmer et al., 2021 adult dataset to the developing inhibitory populations defined in this study. *SCGN*+ INs in the developing brain map to *ADARB2*+ INs in the adult brain.

**Figure S6:**
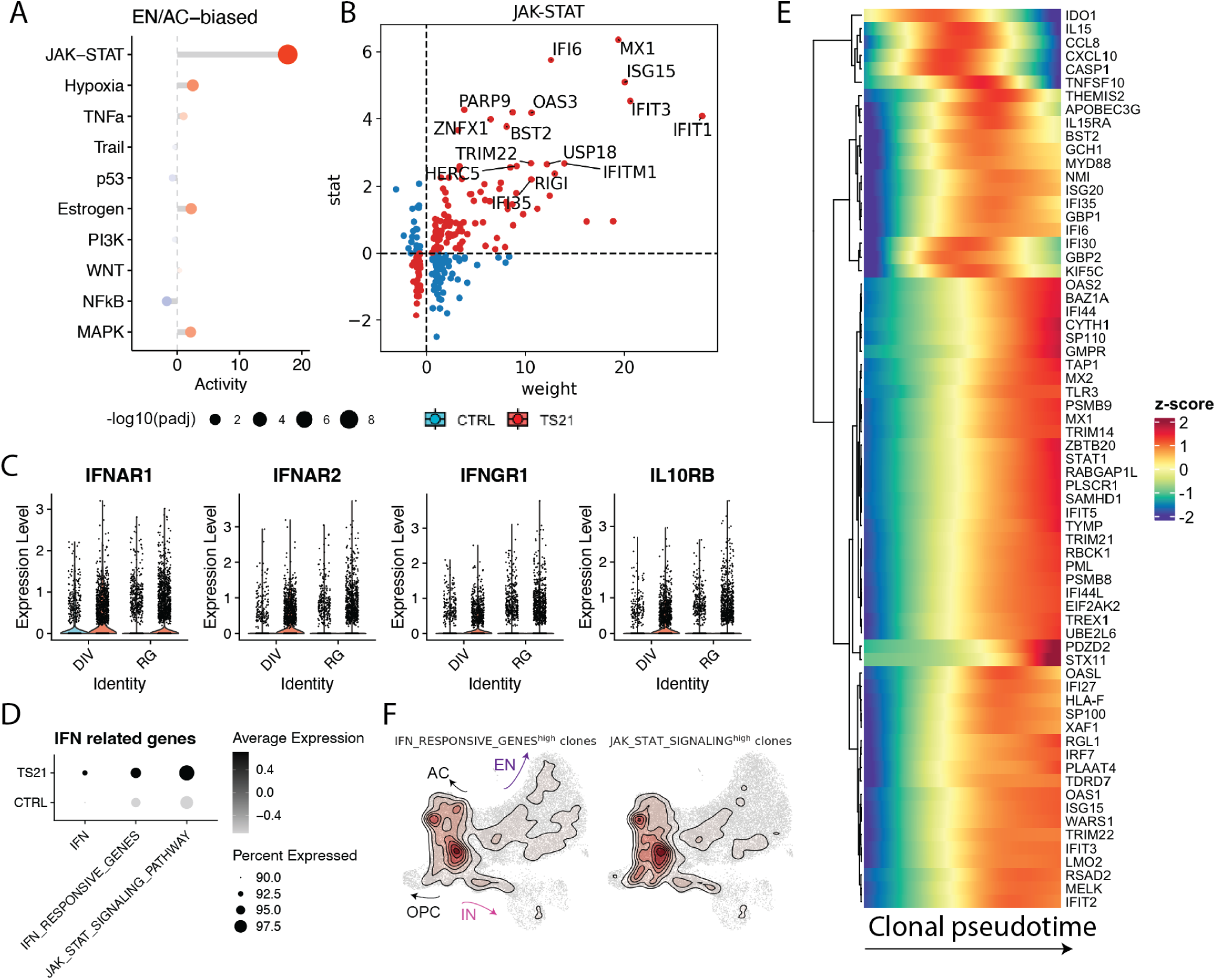
IFN expression in TS21, related to Figure 4. A. Cleveland’s dot plot showing pathway activity in DEGs in EN/AC-biased radial glia class. Size denotes -log_2_(adjusted p value) and color denotes pathway activity calculated using a univariate linear model through PROGENy^54^. B. Scatterplot of genes in JAK-STAT pathway in EN/AC-biased RG. The x-axis denotes the weight of the target gene in the pathway. The y-axis represents the Wald test statistic from DESeq2^35^, defined as the estimated log_2_FC divided by its standard error. C. Violin plot showing expression of four IFN receptor genes on HSA21 in the radial glia class. D. Dot plot showing gene scores of 3 different gene sets downstream of IFN stimulation in the radial glia class, calculated with UCell^56^. E. Heatmap showing gene expression changes of IFN stimulated genes in RG along the clonal pseudotime. Upregulation of these genes at late pseudotime coincides with occurrence of IN-biased clones. F. KDE plots of cells from top 15% clones where the radial glia class has the highest gene score of IFN responsive genes (left) and JAK-STAT signaling pathway (right).

## Supplemental Tables

Table S1. Metadata for samples included in the single cell lineage tracing experiments, related to Figure 1

Table S2. Differentially expressed genes in TS21 by cell type, related to Figure 1

Table S3. Metadata for samples included in the immunohistochemistry experiments, related to Figure 3

Table S4. Differentially expressed genes in TS21 in the radial glia class by clonal cluster, related to Figure 4

## STAR Methods

### EXPERIMENTAL MODEL AND STUDY PARTICIPANT DETAILS

#### Tissue samples

De-identified human tissue samples were collected with previous patient consent in strict observance of the legal and institutional ethical regulations. Protocols were approved by the Human Gamete, Embryo, and Stem Cell Research Committee (institutional review board) at the University of California, San Francisco. Samples used for single cell lineage tracing and immunohistochemical quantification are detailed in Table S1 and S3, respectively.

#### Organotypic slice culture

Tissue samples were embedded in 3% low-melting-point agarose (Fisher, BP165-25) and cut into 300-μm sections perpendicular to the ventricle on a Leica VT1200S vibrating blade microtome in oxygenated artificial cerebrospinal fluid containing 125 mM NaCl, 2.5 mM KCl, 1 mM MgCl2, 1 mM CaCl2 and 1.25 mM NaH2PO4. Slices were cultured in media containing Insulin (Thermo, A1138IJ) -Transferrin (Invitria, 777TRF029-10G) -Selenium (Sigma, S5261-10G), 1.23 mM ascorbic acid (Fujifilm/Wako, 321-44823), 1% polyvinyl alcohol (PVA) (Sigma, P8136-1KG), 100μg/ml primocin (Invivogen, ant-pm-05), Glutamax (Invitrogen, 35050061), 1 mg/mL BSA, 15 μM Uridine (Sigma, U3003-5G), 1 μg/ml L-Glutathione reduced (Sigma, G6013-5G), 1 μg/ml (+)-α-Tocopherol acetate (Sigma, t3001-10g), 0.12 μg/ml Linoleic (Sigma, L1012) and Linolenic acid (Sigma, L2376), 10mg/mL Docosahexaenoic Acid (DHA) (Cayman, 10006865), 5mg/mL Arachidonic Acid (AHA) (Cayman, 90010.1), 20ng/ml BDNF (Alomone Labs, B-250) in DMEM-F12 (Corning, MT10092CM). ROCK inhibitor CEPT cocktail^57^ was supplemented on the first day.

### METHOD DETAILS

#### Plasmids, lentiviral production and transduction

STICR plasmids (Addgene #180483, #186334, #186335) were obtained from the Nowakowski lab. Lentivirus was produced in HEK293T cells seeded at a density of 80,000 cells/cm^2^ 24 hr prior to transfection. Transfection was performed using Lipofectamine 3000 (Invitrogen, L3000001) transfection reagent according to the manufacturer’s protocol. 18 hr post-transfection, the media was replaced and supplemented with 1X ViralBoost (Alstem, VB100). Supernatant was collected 48 hr post transfection and concentrated at 1:100 with lentivirus precipitation solution (Alstem, VC100).

Lentiviral transduction was performed locally at GZ of organotypic slices to preferentially label neural progenitor cells using 1:50-1:100 diluted STICR lentivirus. 24 hr after transduction, virus-containing media was replaced with fresh media and daily half-media exchange was performed. 13-14 days after transduction, slices were dissociated using papain (Worthington, LK003163) supplemented with 5% Trehalose (Fisher Scientific, BP268710), and GFP positive cells were isolated by fluorescence-activated cell sorting (FACS) and captured with PIPseq™ V T10 3ʹ Single Cell RNA Kit and PIPseq™ T20 3ʹ Single Cell RNA Kit v4.0 following the manufacturer-provided protocol (FB0004762; FB0002130).

#### Generation and analysis of scRNA-seq libraries

##### PIP-seq library generation and sequencing

The manufacturer-provided protocol (FB0004762; FB0002130) was used to generate single cell gene expression libraries. To generate STICR barcode libraries, 10 μl of PIP-seq full length cDNA was used as template in a 50 μl PCR reaction containing 25 μl Q5 Hot Start High Fidelity 2X master mix (NEB, M0494) and STICR barcode read 1 and 2 primers (0.5 μM, each) described in Delgado et al.^16^ using the following program: 1, 98 °C, 30 s; 2, 98 °C, 10 s; 3, 62 °C, 20 s; 4, 72 °C, 10 s; 5, repeat steps 2–4 15 times; 6, 72 °C, 2 min; 7, 4 °C, hold. Following PCR amplification, a 0.8–0.6 dual-sided size selection was performed using SPRIselect Bead (Beckman Coulter, B23318). Libraries were sequenced on Illumina NovaSeq platforms to the depth of roughly 25,000 reads/cell for gene expression libraries and 5,000 reads/cell for STICR barcode libraries.

##### Alignments and quality control

PIP-seq^58^ was used to jointly sequence single-cell transcriptomes and clonal barcodes. PIPseeker 3.0 was used to align and trim reads from raw FASTQs jointly from cDNA and STICR libraries. The trimmed reads were then aligned with STARSolo 2.7.11a^59^ using this specific command where $R1 and $R2 are R1 and R2 fastqs from trimmed fastqs generated by PIPseeker, whitelist from PIPseeker, and reference for mouse and human from https://www.thepoollab.org/resources:

STAR --genomeDir ∼/human_GRCh38_optimized_reference_v2_STAR --runThreadN 16 --soloType CB_UMI_Simple --soloCBstart 1 --soloCBlen 16 --soloUMIstart 17 --outSAMattributes CB CR CY GX GN UB UR UY NH HI nM AS sF --outSAMtype BAM SortedByCoordinate --soloCBmatchWLtype Exact --soloUMIdedup 1MM_CR --soloFeatures Gene SJ GeneFull GeneFull_Ex50pAS GeneFull_ExonOverIntron Velocyto --soloMultiMappers EM --soloCellReadStats Standard --soloCellFilter EmptyDrops_CR --soloUMIfiltering MultiGeneUMI_CR --outSAMunmapped Within --soloBarcodeReadLength 0 --readFilesCommand zcat --limitBAMsortRAM 1775716961230000 --soloCBwhitelist barcodes/barcode_whitelist.txt --soloUMIlen 12 --readFilesIn $R2 $R1

Ambient RNA was filtered using FastCAR^60^ and cells were filtered for a minimum of 500 genes, mitochondrial cutoff at 10% of total transcripts, and doublet score of less than 1 using scds^61^.

##### STICR barcode assignments

STICR barcodes were aligned and assigned using a modified NextClone^62^ workflow that allows for barcode whitelisting. The pipeline is available at: https://github.com/cnk113/NextClone. Individual barcodes were filtered by at least 2 reads supporting a single UMI and at least 2 UMI to call cells with a barcode. Clone calling was done using CloneDetective^62^. Cells derived from multicellular clones that have a minimum of 3 cells were included for clonal analysis.

##### Reference mapping and cell type annotation

The developing human cortex multiomic dataset was obtained from Wang et al.^17^ and used for reference mapping to support cell type annotation in this study. The reference model was built with scvi-tools^63^ using top 2500 variable genes defined in the *in vivo* dataset and used for integration and label-transfer to the *ex vivo* query dataset generated in this study. Cell type annotation was then performed based on marker expression as well as predictions from scANVI. Pearson correlation coefficients between the two datasets were calculated using top 25 markers in each cell type identified from the *in vivo* dataset to examine the fidelity of cell identities collected in this study.

##### Clonal analysis

Clones with less than three cells from the RG lineage were removed from the clonal analysis. Cospar^27^ was used to calculate the fate coupling scores, defined as the normalized barcode covariance between different cell types. scLiTr^52^ was used to construct the clonal embedding space (clone2vec) and identify clusters of clones with distinct fate biases by training a neural network to infer clonal labels of nearest neighbors for each clonally annotated cell. To minimize batch effects in cluster identification, we used integrated data from all 14 individuals to define six clusters with distinct fate biases.

To examine stage-dependent changes between CTRL and TS21, samples were binned into two age groups: early (GW14-17) and late (GW18-GW23). Milo^31^ was then used for cluster-free differential abundance testing as detailed below. Monocle3^53^ was used to construct a pseudotime trajectory at the clone level on the UMAP output by scLiTr. Pseudotime distribution in each stage between conditions was visualized to examine the changes of RG developmental tempo. The radial glia class (RG, DIV) from multicellular clones was subset and assigned with clonal pseudotime values to fit gene expression changes along the clonal pseudotime.

To investigate the gene expression differences in RG of different fate biases, differential expression analysis was performed following the DEseq2^35^ pipeline in the subset radial glia class to contrast RG from each clonal cluster between conditions as detailed below. Pathway scoring was performed using PROGENy^54^ to identify enriched pathways in each fate biased RG cluster.

##### Differential composition and gene expression analysis

Milo^31^ was used to test differential abundance at the cell level and at the clone level. Briefly, clones or cells from multicellular clones were grouped into four groups by disease status and age ‘status_group’: CTRL_early, CTRL_late, TS21_early, TS21_late. Cells from each ‘stage_group’ were randomly subset to 2,000 cells per group to ensure the balance of total cell numbers between groups. Neighborhoods were defined using KNN graphs computed in the latent space generated by clone2vec and SCANVI for clones and cells, respectively. Differential abundance in each neighborhood was tested using design = ∼ 0 + stage_group. All pairwise contrasts were computed, and log fold changes in neighborhoods with significance were visualized.

Cluster-aware differential gene expression analysis was performed using DEseq2^35^ contrasting CTRL and TS21 using age range as covariate. When pseudobulking within clusters, conditions that have less than 10 cells per cluster or less than 2 biological replicates were removed from downstream analysis. The same pipeline was applied at the clone level between clonal clusters in the radial glia class and at the cell level between cell types in all cells.

#### Somatic lineage tracing analysis

10x short-read libraries of the adult PFC dataset obtained from Palmer et al., 2021^12^ were processed and aligned using STARSolo. Paired PacBio Iso-seq of single-cell libraries were initially processed with flexiplex^64^ to identify cell barcode and UMI regions in the fastq and then aligned with ultra to the cellranger reference transcriptome. A script was used to add CB/UMI to the aligned long-read BAMs and then using subset-bam from 10X Genomics, the BAM files were split into pseudobulked cell-type BAMs. Variants were called at cell-type resolution using longcallR^65^ to identify variants on cell-type BAMs and filtered for RNA editing sites. After filtering VCFs, single-cell genotyping was performed on the short-read libraries using VarTrix from 10X Genomics. The resulting raw somatic cell-by-variant matrix was then pseudobulked at cell-type resolution for Pearson correlation and then normalized to somatic variants in DL ENs.

Reference mapping was performed to link developmental IN subtypes generated in this study to their adult counterparts. Briefly, short-read libraries from Palmer et al., 2021 were used to build a reference model using top 2500 highly variable genes using scvi-tools. Label transfer to the cells generated in this study was then performed using SCANVI to examine correspondence of these cell types.

#### Immunohistochemistry

Primary cortical tissue were fixed with 4% paraformaldehyde (PFA) in PBS overnight, washed three times with PBS, then placed in 10% and 30% sucrose in PBS overnight, sequentially, and embedded in OCT for sectioning to 20 μm.

All samples were blocked with blocking solution (5% BSA, 0.3% Triton-X in PBS) for 1 hr. Primary and secondary antibodies were diluted in blocking solution. Samples were incubated in primary antibody solution overnight at 4C, then washed three times with PBS at room temperature. Samples were then incubated in a secondary antibody solution with DAPI for 1 hr at room temperature and then washed three times with PBS before mounting samples on slides with Fluoromount (Invitrogen, 0100-20). Primary antibodies used in this study include rabbit-NEUROD2 (1:500, Abcam, ab104430) and goat-SCGN (1:500, R&D, AF4878). Secondary antibodies in this study include donkey anti rabbit 555 (1:1000, Thermo, A11012) and donkey anti goat 488 (1:1000, Thermo, A11055). Images were collected using 20x air objectives on an Evos M7000 microscope, and processed using ImageJ/Fiji.

#### Image quantification

Individuals used for immunohistochemistry and quantification are detailed in Table S3. Images shown in figures were representative of multiple images taken across multiple replicates, and tile scans for given stainings are shown in supplemental images.

To minimize technical biases, we developed an automated scalable pipeline using Cellpose^32^ for signal segmentation and quantification. Briefly, regions of interest (ROIs) for GZ(SVZ) and IZ/SP were cropped using macros in ImageJ/Fiji. Unprocessed images were fed into Cellpose with the following parameters: model = cyto3; diameter = 30 pixels or none; flow threshold = range between 0.1 - 0.6; cellprob threshold = range between 0 - 3; restoration = upsample_cyto3 or deblur_cyto3 or none. Quality control check was performed by overlaying segmentation masks onto the original images.

Quantification of marker positive populations in each individual was done by summing the overlap of DAPI and marker positive cell counts over DAPI counts across 4-8 ROIs from each tile scan and averaging across 2-4 tile scans. Statistics for *in vivo* comparison between CTRL and TS21 in Figure 3 were performed on a total of 50 tile scans taken from 21 independent biological replicates, aged GW14-GW23.5. Two-sided Wilcoxon test was performed after binning samples into two age groups (early, GW14-18; late, GW19-23.5).

### QUANTIFICATION AND STATISTICAL ANALYSIS

Statistical analyses are described in the figure legends with additional details in the corresponding METHOD DETAILS sections. The n values can be found in the main text and/or figure legends. Individuals included for the single cell lineage tracing in Figure 1 and immunohistochemical quantification in Figure 3 are detailed in Tables S1 and S3, respectively.

## Notes

### Competing Interest Statement

The authors have declared no competing interest.

